# Geometric Tuning of Cytokine Receptor Association Modulates Synthetic Agonist Signaling

**DOI:** 10.1101/2025.10.12.681819

**Authors:** Marc Expòsit, Mohamad Abedi, Aditya Krishnakumar, Shruti Jain, Ta-Yi Yu, Timothy R. Hercus, Divĳ Mathew, Sophie Gray-Gaillard, Zhĳie Chen, William S. Grubbe, Andrew Favor, Winnie L. Kan, Thomas Schlichthaerle, Wei Chen, Michael W. Parker, Juan L. Mendoza, Angel F. Lopez, E. John Wherry, David Baker

## Abstract

Cytokines signal by bringing receptor subunits together, but the role of receptor geometry in shaping signaling remains unclear because natural ligands enforce fixed assemblies. Here, we present a *de novo* protein design platform that rigidly scaffolds receptor-binding domains into defined spatial arrangements. Applying this across IL-7, type I and III interferons, IL-10, gp130, β common, and synthetic receptor pairs, we show that by varying geometry, we can bias pSTAT pathway usage and tune functional outcomes. Geometric control allowed us to decouple pSTAT1 from pSTAT5 in IL-7, separate antigen presentation (MHC-I) from checkpoint induction (PD-L1) in type I interferons, and suppress pro-inflammatory IFNγ secretion while retaining anti-inflammatory activity in IL-10. We further created minimal IL-6 and IL-3 agonists and strengthened synthetic receptor pairings inaccessible with present cytokines. These results establish receptor geometry as a central determinant of cytokine activity and provide a platform for programmable immune modulation.

## Introduction

Cytokines drive immune responses by assembling receptor subunits to activate intracellular signaling cascades, primarily through the JAK/STAT pathway^1,2^, but their therapeutic use is constrained by pleiotropy and systemic toxicity^3–5^. Engineering efforts have largely focused on modulating receptor-binding affinity^6,7^, leaving underexplored the role of receptor geometry, a variable that could strongly influence signaling efficiency and specificity^8^. Existing cytokines impose fixed dimerization geometries, making this axis difficult to probe, though structural studies suggest that even subtle geometric shifts can have functional consequences^9^. Prior work with engineered erythropoietin and NGF ligands demonstrated that altering receptor spacing in homodimers can redirect signaling, but this has not been extended to heterodimeric cytokines or generalized across receptor families^10–12^. Thus, the fundamental relationship between receptor geometry and signaling outcomes remains unclear.

We reasoned that the relationship between receptor geometry and signaling outcome could be systematically explored for a range of cytokine-receptor systems by using protein design to create *de novo* designed signaling molecules (Novokines), which rigidly pair designed receptor binding proteins in a range of orientations and spacings, systematically varying receptor spacing and composition (**Figure 1A**). We set out to use this approach to map how geometry shapes signaling across natural and synthetic receptor pairs, and to generate biased agonists with therapeutic potential.

**Figure 1:**
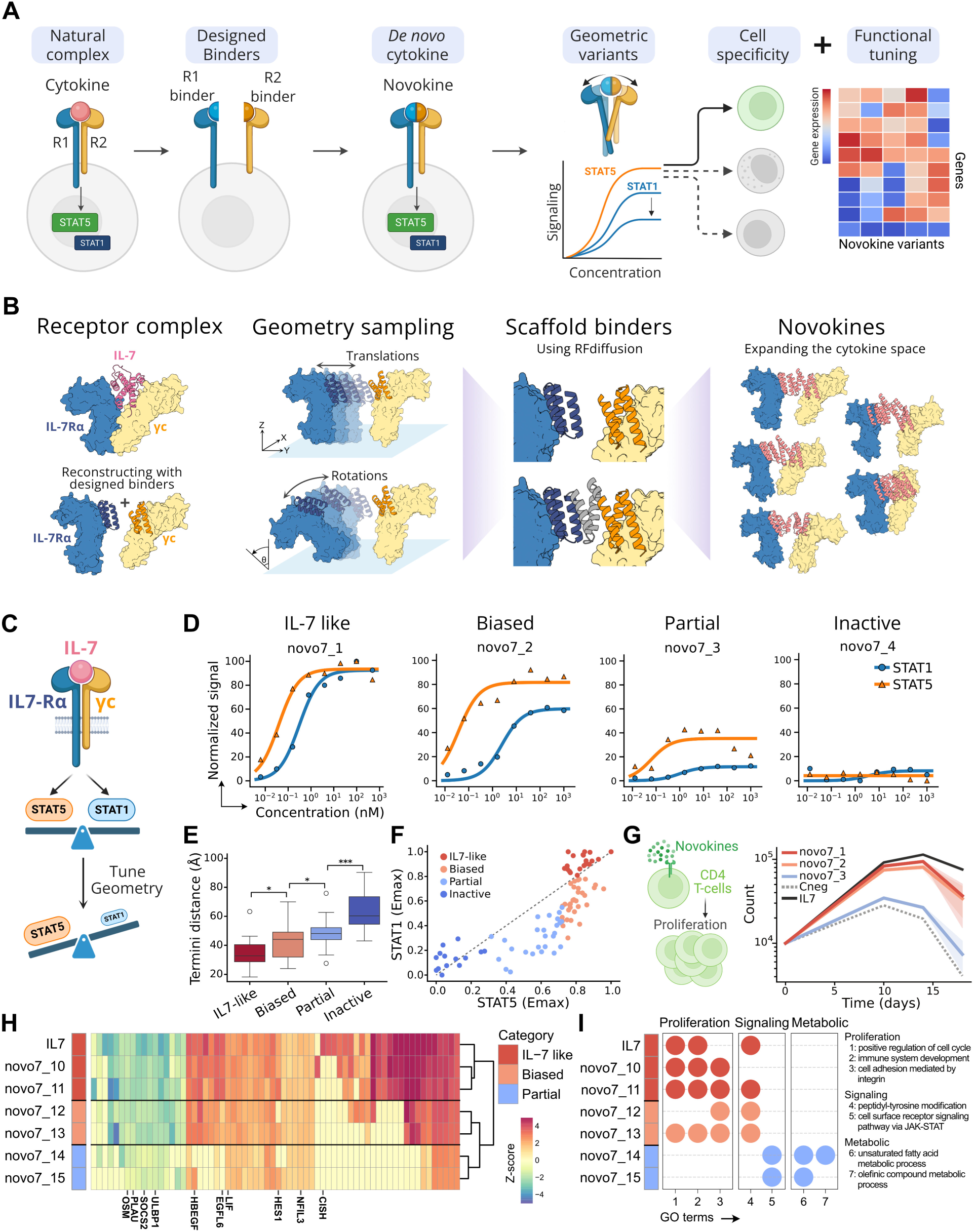
Design of *de novo* IL-7-like cytokines with geometrically tuned signaling. (A) High level representation of our generalizable platform tuning cytokine receptor geometry to alter pSTAT signaling and functional outcomes. (B) Method for design of *de novo* agonists, using IL-7 as an example. The structure of the IL-7 signaling complex is used as a reference but recreated with binders for IL-7Rα and γc. The geometry of the complex is altered by sampling any arbitrary set of translations and rotations for one of the two receptors and designing scaffolds for the binders that rigidly stabilize the receptor positions. The resulting IL-7-like cytokines sample a wide range of receptor geometries. (C) Diagram on IL-7 signaling. Tuning geometry aims to decouple the balance between pSTAT1 and pSTAT5 elicited by the natural cytokine. (D) pSTAT1 and pSTAT5 signaling of four selected geometry variants, displaying IL-7 like (strong in both pSTATs), biased (weaker pSTAT1), partial (weaker for both pSTATs) or no signaling. (E) Distances between the membrane-proximal receptor termini classifying the designs by signaling profile. Adjusted *p* values as: ns: *p* > 0.05; *: *p* ≤ 0.05; **: *p* ≤ 0.01; ***: *p* ≤ 0.001. (F) Comparison of the Emax in pSTAT1 and pSTAT5 between the set of geometrically tuned agonists. (G) CD4^+^ T-cell proliferation over 18 days. (H) Clustering designs by their transcriptional response in CD4^+^ T-cells assessed by RNA-seq, showing the set of top 30 differentially expressed genes per design. Gene symbols relevant to IL-7 downstream signaling have been included. (I) Gene set enrichment analysis presenting relevant GO terms identified in the differentially expressed genes for each design.

## Results

### De Novo Design of IL-7 and Functional Tuning via Receptor Clustering Geometries

We started with IL-7 as a model system to explore how receptor geometry encodes cytokine function. IL-7 plays an essential role in T-cell development and homeostasis by activating pSTAT5 and pSTAT1, but has not previously been engineered^13,14^. IL-7 naturally signals by dimerizing IL-7Rα with the common γ chain (γc), bringing the intracellularly associated JAK1 and JAK3 into proximity to initiate phosphorylation cascades^15,16^. By rebuilding IL-7 from first principles we aimed to systematically vary IL-7Rα-γc clustering geometries, generate partial agonists with distinct transcriptional programs and reveal how receptor assembly governs signaling outcomes (**Figure 1B**).

To reconstitute IL-7 signaling, we used previously designed minibinders (∼60 amino acids) that independently recognize IL-7Rα and γc^17,18^. By fusing these binders with flexible glycine-serine linkers, we generated IL-7 mimics that engage both receptors and successfully reconstitute IL-7 signaling. However, varying linker length or fusion orientation did not appreciably tune signaling as observed through the ratio of pSTAT1 to pSTAT5 activation (**Figure S1**). To systematically probe the role of receptor geometry, we next moved beyond flexible linkers to create rigid geometric fusions by computationally sampling thousands of arbitrary rotations and translations of IL-7Rα and γc around their natural architecture in the signaling competent complex. Using RFdiffusion, we scaffolded the binders into rigid architectures that preserved each predefined geometry, thereby stabilizing the receptors in diverse spatial configurations (**Figure 1C**). From this set, we selected a panel of 96 geometrically distinct candidates, novo7, for functional testing in primary human peripheral blood mononuclear cells (PBMCs).

Functional characterization of the novo7 panel revealed four distinct signaling phenotypes (**Figure 1D**). Some designs fully recapitulated wild-type IL-7 activity, strongly activating both pSTAT5 and pSTAT1; others exhibited biased agonism, maintaining pSTAT5 signaling while attenuating pSTAT1. A third group displayed partial agonism, with reduced activation of both STATs, while a final class was inactive. Across the panel, signaling activity correlated with the distance between the membrane-proximal termini of IL-7Rα and γc and there was no correlation with binding affinities across designs that explain these differences (**Figures 1E, S2, and S3**). pSTAT1 signaling was particularly sensitive to this spacing: biased agonists consistently showed sharper reductions in pSTAT1 activation, while pSTAT5 declined more gradually, suggesting stricter geometric requirements for pSTAT1 (**Figure 1F**). To test whether these signaling patterns translated into functional consequences, we measured CD4+ T-cell proliferation over 18 days. IL-7-like and pSTAT5-biased agonists supported robust proliferation, whereas partial agonists failed to do so (**Figure 1G**).

To evaluate whether these functional differences could be predicted from transcriptional outcomes, we performed RNA sequencing (RNA-seq) on CD4+ T cells stimulated with IL-7-like, biased, and partial agonists. Each class induced a distinct gene expression program: IL-7-like agonists clustered closely with natural IL-7, biased agonists formed a separate group, and partial agonists diverged most strongly (**Figures 1H, S4, and S5)**. Gene set enrichment analysis revealed that only IL-7-like and biased agonists upregulated pathways associated with proliferation, explaining why partial agonists failed to support proliferation **(Figure 1I**). Together, the results demonstrate that receptor geometry can be precisely engineered to decouple distinct aspects of cytokine signaling and that these differences propagate through gene expression programs to produce divergent cellular behaviors.

### Geometric Tuning of Type I Interferon Decouples Antigen display from Checkpoint inhibition

Type I interferons (IFNs) coordinate host immunity across viral infection and cancer, yet their simultaneous induction of immune-activating and immune-suppressive programs creates a fundamental duality in their biological effects^19,20^. Two IFN-stimulated genes illustrate this tension: MHC class I, which promotes antigen presentation to cytotoxic T cells, and PD-L1, an immune checkpoint ligand that suppresses effector T-cell activity through PD-1 engagement^21^. In both contexts, these genes are typically co-induced, coupling immune recognition with immune inhibition and potentially limiting therapeutic benefit^22^. The ability to decouple MHC-I and PD-L1 regulation could enable more precise immune modulation, boosting tumor recognition while minimizing immune suppression^23^. We therefore asked whether geometric tuning of receptor association could uncouple these outputs (**Figure 2A**).

**Figure 2:**
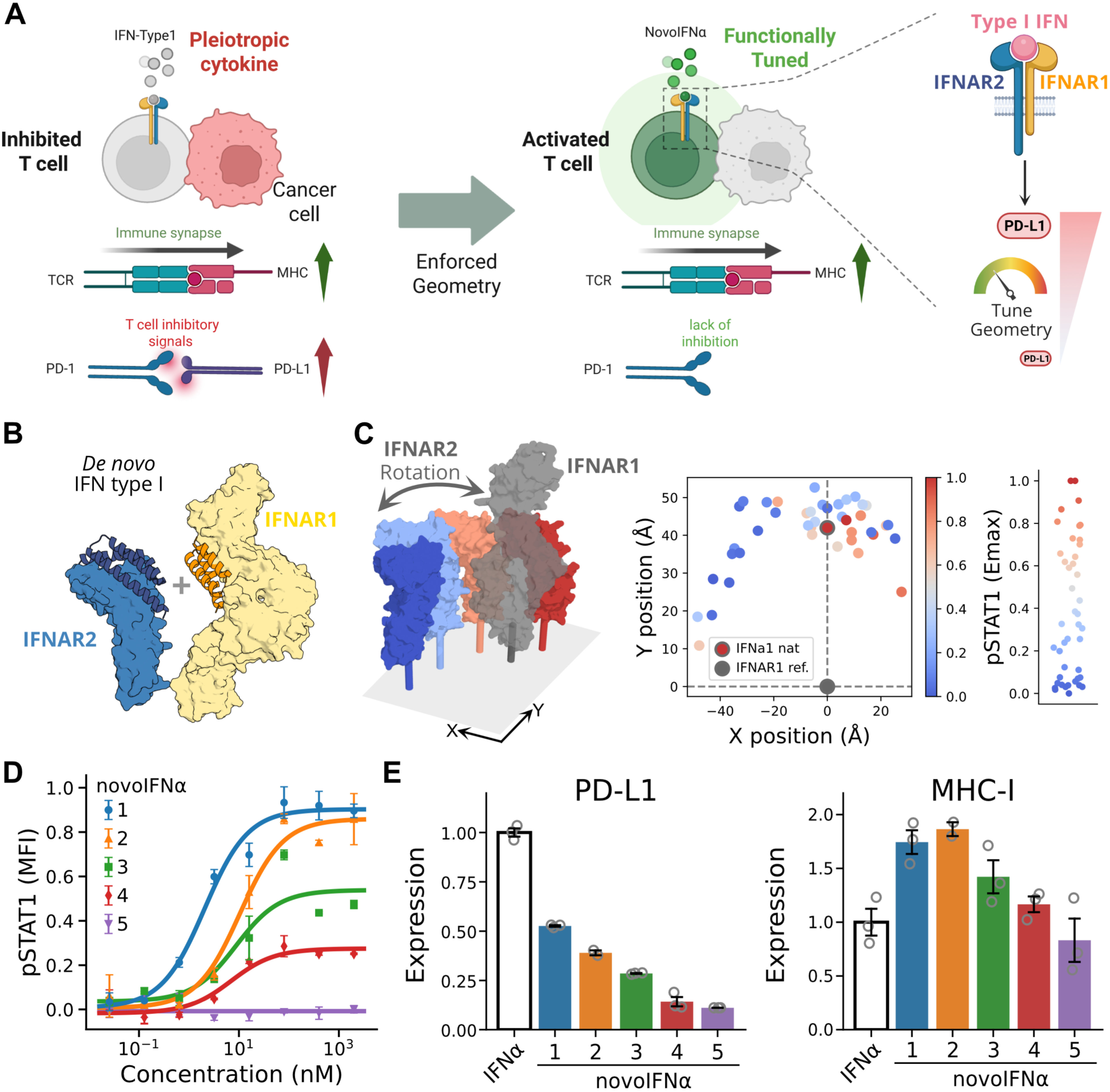
Geometrically engineered type I interferon cytokines enhance anti-tumor responses. (A) Diagram comparing the activities of natural type I IFNs, which upregulate both immune-activating and suppressor signals resulting in T-cell inhibition, and the *de novo* ligands novoIFNα, which promote T-cell activation by reducing the overexpression of immunosuppressive signals while maintaining immune-activating ones. (B) Creating structurally diverse type I IFN-like novokines, novoIFNα, by combining binders for IFNAR1 and IFNAR2. (C) Aligning the IFNAR1 chain of all designed agonists and representing the position of the membrane-proximal domain of the reference IFNAR1 (gray) and the corresponding IFNAR2, depending on the geometry of the agonist (colored by Emax), including the natural IFNa1 ligand (gray edged dot). (D) pSTAT1 dose-response curve of selected novoIFNα agonists. (E) Expression of PD-L1 (left) and MHC-I (right) by the natural IFNa1 or designed novoIFNαs in a tumor cell line.

Type I IFNs are particularly suited for this question, as they comprise more than 13 natural isoforms that all signal through the same receptor pair, IFNAR1 and IFNAR2, yet elicit distinct transcriptional responses^24^. While isoform diversity has been attributed to differences in ligand-receptor binding affinity, the contribution of receptor geometry remains largely unexplored^25–27^. Prior efforts to alter geometry, such as inserting residues into transmembrane domains to rotate intracellular domains, were coarse (∼100° per insertion) and failed to reveal consistent effects, while more recent work using flexible extracellular VHH fusions explored only a narrow range of geometries and produced biased agonists^28,29^. To directly investigate the role of geometry, we generated a library of 96 synthetic type I IFN agonists (novoIFNα) by rigidly fusing IFNAR1- and IFNAR2-specific binders across a wide range of predefined spatial arrangements (**Figure 2B**). These designs stabilized the receptor dimer in geometries independent of ligand affinity and produced a spectrum of STAT1 phosphorylation responses in PBMCs, ranging from wild-type-like activation to partial agonism or inactivity (**Figures 2C, 2D, and S6**). Unlike IL-7, where distance was the primary geometric factor influencing signaling, type I IFN activity was strongly impaired by lateral deviations from the native complex of approximately ∼90°, highlighting unique geometric sensitivities of this signaling system (**Figure 2C**).

To assess functional outcomes, we measured MHC-I and PD-L1 expression in a cancer cell line one day after stimulation. Whereas wild-type IFN co-induced both markers, several of the novoIFNα agonists maintained or enhanced MHC-I expression with reduced PD-L1 induction (**Figure 2E**). The reduction in PD-L1 induction strongly correlated with pSTAT1 intensity (R² = 0.54, Spearman’s ρ = 0.74, p<0.005, permutation test), linking geometrically induced pSTAT1 damping to functional changes in PD-L1 expression (**Figure S7**). Thus, by selectively tuning receptor geometry, we generated biased type I IFN agonists that promote immune activation while minimizing immunosuppressive feedback, offering a strategy to enhance anti-tumor responses.

### Geometric Tuning of IL-10 modulates Cell-Specific Anti- and Pro-inflammatory Activity

We next focused on IL-10, a major anti-inflammatory cytokine with therapeutic potential in cancer and autoimmune disorders such as inflammatory bowel disease^30^. Clinical translation of IL-10 has been limited by its dual activity: while it suppresses immune responses by inhibiting monocyte and macrophage activation, it simultaneously can promote pro-inflammatory cytokine secretion, notably IFN-γ from CD8+ T cells^31^. Prior strategies to bias IL-10 function have included reducing receptor affinity to dampen CD8+ T-cell activation or enhancing local delivery via engineered bacteria^32,33^. Given our success in generating biased and partial agonists of IL-7 and type I IFNs, we hypothesized that geometric tuning of the IL-10 receptor complex could similarly decouple these opposing activities and generate more selective anti-inflammatory function (**Figure 3A**).

**Figure 3:**
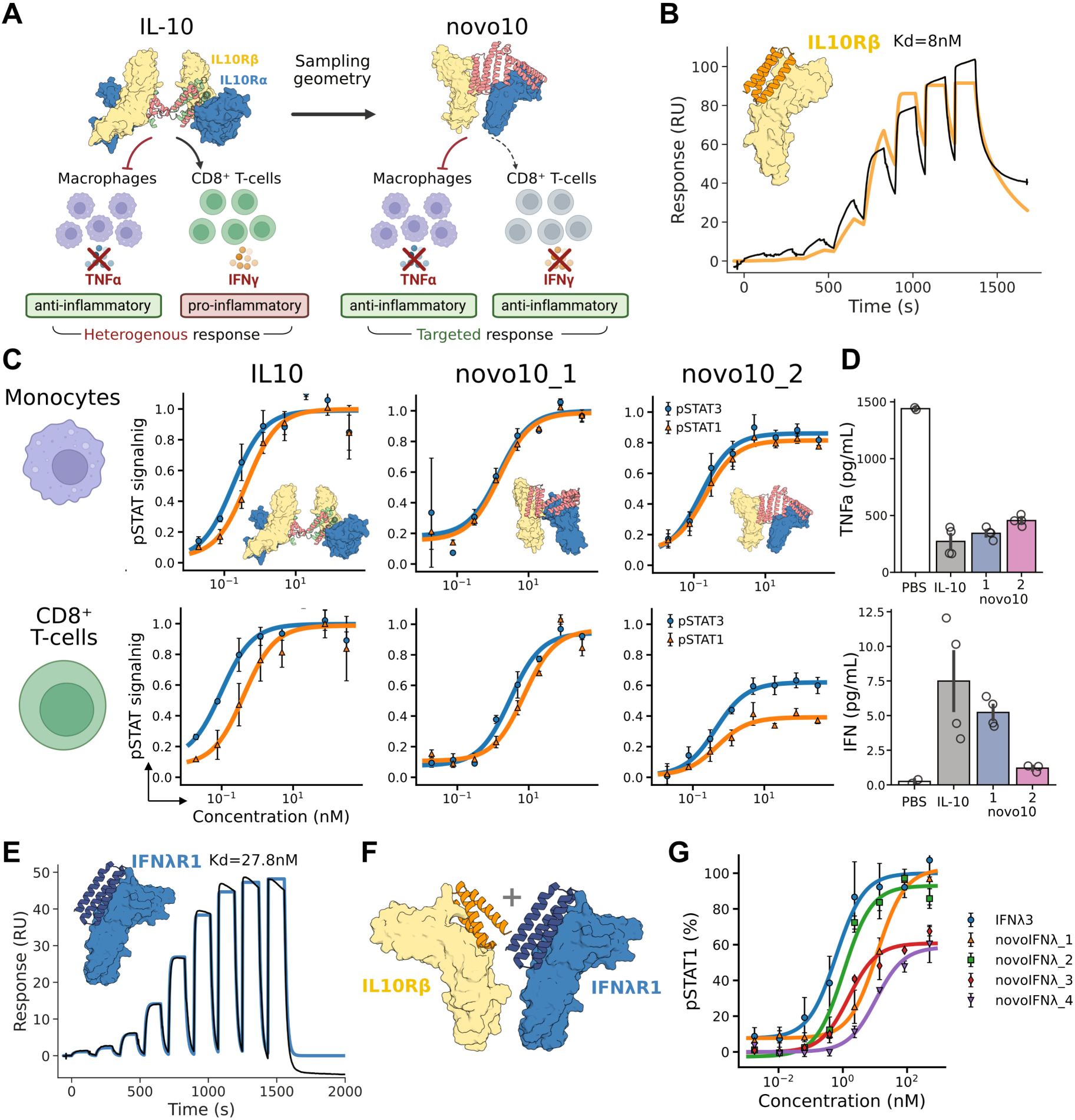
Engineered IL-10 variants with suppressed pro-inflammatory response and geometric tuning of type III interferon signaling. (A) Schematic representation of the heterogeneous effects of natural IL-10 in PBMC subsets. While IL-10 prevents monocytes and macrophages from secreting TNFα when challenged, eliciting an anti-inflammatory effect, it also induces secretion of IFN-γ by CD8^+^ T cells, which is pro-inflammatory. Geometrically tuned variants aim to attenuate the secretion of IFN-γ by CD8^+^ T cells, removing the pro-inflammatory signature and achieving a targeted anti-inflammatory effect. (B) Design and characterization of an IL-10Rβ binder. (C) pSTAT signaling (pSTAT1 in orange, pSTAT3 in blue) of natural IL-10 compared to the novokines novo10_1 and novo10_2, profiled in monocytes (top) and CD8^+^ T cells (bottom) subpopulations of PBMCs. (D) Secretion of TNFα (top) or IFN-γ (bottom) when testing selected ligands in bulk PBMCs challenged with LPS or isolated CD8^+^ T cells, respectively. (E) Design and characterization of an IFN-λR1 binder. (F) Exploring geometry in type III IFN by combining binders for IFN-λR1 and IL-10Rβ in different orientations. (G) pSTAT1 signaling in the A549 epithelial cell line of designed ligands compared to natural type III IFN (IFNλ3).

Native IL-10 is a domain-swapped homodimer that engages a tetrameric receptor complex consisting of two IL-10Rα and two IL-10Rβ chains. Because monomeric IL-10 variants that recruit only one IL-10Rα/IL-10Rβ pair can still signal^34,35^, we constructed synthetic IL-10 agonists by rigidly scaffolding *de novo* binders for IL-10Rα and IL-10Rβ into ligands that stabilize dimeric, but not tetrameric, assemblies. To enhance the affinity of the previously described IL-10Rβ binder, we used partial diffusion to computationally enhance its affinity from 200nM to 8nM (**Figure 3B**). Testing these designs in PBMCs revealed distinct signaling behaviors: one variant (novo10_1) fully recapitulated wild-type IL-10 activity in both monocytes and CD8+ T cells, while another (novo10_2) exhibited cell-type-specific biased agonism. Novo10_2 maintained strong pSTAT1/3 signaling in monocytes but reduced pSTAT1/3 activation in CD8+ T cells, with pSTAT1 being more strongly reduced than pSTAT3 (**Figure 3C**). These results demonstrate that dimeric assemblies are sufficient for IL-10-like activity, and that geometric tuning can shift signaling in a cell-type-specific manner.

To investigate whether the reduced pSTAT1/3 ratio in CD8+ T cells attenuated pro-inflammatory outputs, we measured cytokine secretion following stimulation. While wild-type IL-10, novo10_1 and novo10_2 all suppressed TNFα release from monocytes upon bacterial lipopolysaccharide (LPS) challenge, only the biased agonist novo10_2 markedly reduced IFN-γ secretion from activated CD8+ T cells (**Figure 3D**). This agrees with the observed reduction in pSTAT1 signaling, a key driver of IL-10-mediated IFN-γ induction, and confirms that geometric tuning can selectively modulate pSTAT1/3 activation in CD8^+^ T cells to eliminate pro-inflammatory activity, while preserving anti-inflammatory suppression.

We next expanded to type III IFNs, which signal through dimerization of IFN-λR1 and IL-10Rβ, a receptor complex structurally similar to type I IFNs but intrinsically less active^36^. Type III IFNs (IFN-λs) are attractive therapeutic candidates because they elicit antiviral responses similar to type I IFNs but with greater tissue specificity and reduced systemic toxicity^37,38^. Prior work showed that rotating the IFN-λR1 intracellular domain (ICD) by alanine insertions in the transmembrane region modified pSTAT1 activation^39^, but this approach is not clinically translatable. Also, recently, it was shown that IFN-λ4 naturally engages the receptors with a different geometry than the canonical type III IFN, IFN-λ3^9^. We therefore asked whether extracellular geometry alone could recapitulate these effects.

Using RFdiffusion binder design, we first created an IFN-λR1 binder with a 27.8 nM affinity, which was then fused with the IL-10Rβ binder to generate synthetic type III IFN novokines (**Figure 3E**). The resulting novokines, novoIFNλ, spanned diverse geometries, including those mimicking the rotations of the intracellular domains produced by alanine insertions. When evaluating their activity in an epithelial cell line, several designs activated pSTAT1 with Emax values comparable to wild-type IFN-λ, and some decreased the maximum response (**Figures 3F and 3G**). This suggests that extracellular geometry can modulate signaling like receptor alanine insertion, with the advantage that the ligand, not the receptor, is engineered, enabling potential protein therapeutic approaches downstream.

### Strengthening and Modulating Novel Cytokines by tuning their clustering geometries

The modularity of our design framework allows us to move beyond natural receptor pairings and explore combinations for which no cytokine exists^29,40,41^. As shown in previous work, flexibly linking *de novo* designed receptor binders can generate diverse synthetic ligands that engage non-natural receptor pairs and elicit novel cellular responses, although many of these constructs signal weakly^18^. Because geometric tuning allows us to span a wide range of signaling strengths, we reasoned that enforcing receptor geometry might enhance signaling by stabilizing receptor orientation within these novel assemblies that were not optimized by nature.

We focused on three synthetic ligands identified in the companion study that signaled weakly: IL-2Rβ/IFNAR1, IL-2Rβ/IL-10Rβ, and IL-10Rα/γc. The first pair was selected because the companion study showed IFNAR1 can signal robustly with multiple cytokine receptors, similar to common receptors like γc or βc^18^. The other two were highlighted in previous studies, with the IL-10Rα/γc intracellular domain combination recently demonstrated to have potent anti-tumor effects^29,41^. A likely explanation for the weak signaling observed with flexible binder fusions is that native receptor pairs benefit from stabilizing transmembrane domain interactions, which might be absent in synthetic combinations^42^. We therefore hypothesized that rigid fusion of *de novo* receptor binders could compensate by stabilizing extracellular domain geometry. Indeed, rigid novokines markedly enhanced pSTAT5 signaling relative to flexible linker designs in either orientation, suggesting that geometric stabilization can restore functional signaling even in pairs not naturally evolved for transmembrane interactions (**Figure 4A**). Notably, this approach would enable induction of anti-tumor activity by IL-10Rα/γc in a physiological context using a protein therapeutic, avoiding the need for engineered cell therapies.

**Figure 4:**
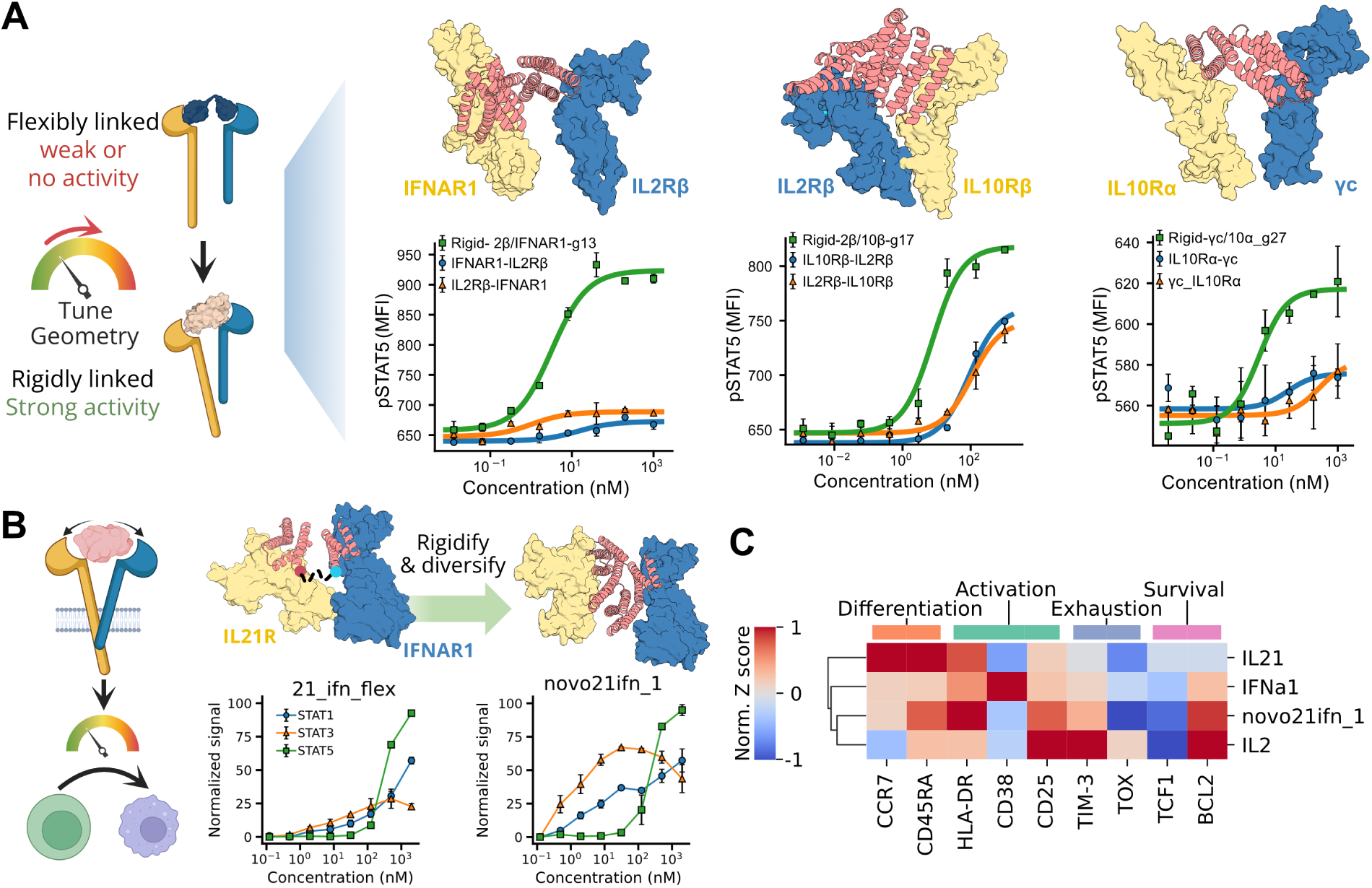
Enhancing and modulating signaling of non-natural receptor pairs through geometric changes. (A) Rigid fusions of binders engaging non-natural cytokine pairs enhance signaling compared to flexible fusions. Flexible fusions are labeled according to the N- to C-terminal order in which the binders are fused. (B) Signaling profiles of a flexible fusion between IL-21Rα and IFNAR1 binders and their tunability when using rigid fusions. (C) Expression profiles of differentiation, activation, exhaustion, and survival markers in naive CD8^+^ T cells treated with anti-CD38/anti-CD3 beads, supplemented with natural cytokines or designed novokines for 4 days.

We next examined a newly identified novokine engaging IL-21Rα and IFNAR1, a receptor pair with no structural precedent^18^. To explore the signaling space of this pair, we sampled 24 geometries that brought the membrane-proximal domains into proximity while avoiding steric clashes. Rigidly fused IL-21Rα/IFNAR1 designs showed more than threefold increases in pSTAT1/3 signaling potency, and one variant (novo21ifn_1) also boosted pSTAT5 activity (**Figure 4B**). To evaluate functional consequences, we profiled cell surface markers on CD8+ T-cells using flow cytometry after CD3/CD28 stimulation and treatment with designed or natural cytokines, as IL-21 is a key regulator of T-cell specialization and maintenance. The biased novo21ifn_1 rigid ligand induced a profile closer to IL-2, with increased BCL2 expression supporting survival and reduced TCF1 levels, consistent with effector differentiation, in contrast to the profiles seen with IFNa1 and IL-21 (**Figures 4C and S8**). The rigid novokine novo21ifn_1 further enhanced activation (HLA-DR) and reduced exhaustion (TOX, TIM-3) compared to IL-2, making it a promising candidate for promoting T-cell activation and persistence. These findings suggest that geometric tuning of novel receptor pairs enables functional rewiring of cytokine activity.

Together, these results establish a new paradigm in cytokine engineering: by identifying receptor pairs enriched in a target cell type and sampling receptor geometry, it is possible to create novel cytokines with tunable activities, expanding the functional repertoire of immune signaling beyond what is encoded in nature.

### Designing Minimal and Geometry-Constrained Cytokine Agonists

To test the broader applicability of our framework, we extended it to IL-6 and IL-3, two cytokines with complex multimeric receptor systems. IL-6 signals through a hexameric complex of IL-6R and gp130, but only gp130 is required for downstream activity^43–46^. We designed minimal agonists by directly engaging gp130 using homodimeric fusions assembled with the WORMS framework (**Figure 5A**)^47^. One of these designs (novo6_1) recapitulated native pSTAT1/3 activity, while activating pSTAT5, indicating that small geometric shifts in gp130 arrangement can redirect signaling (**Figure 5B**).

**Figure 5.**
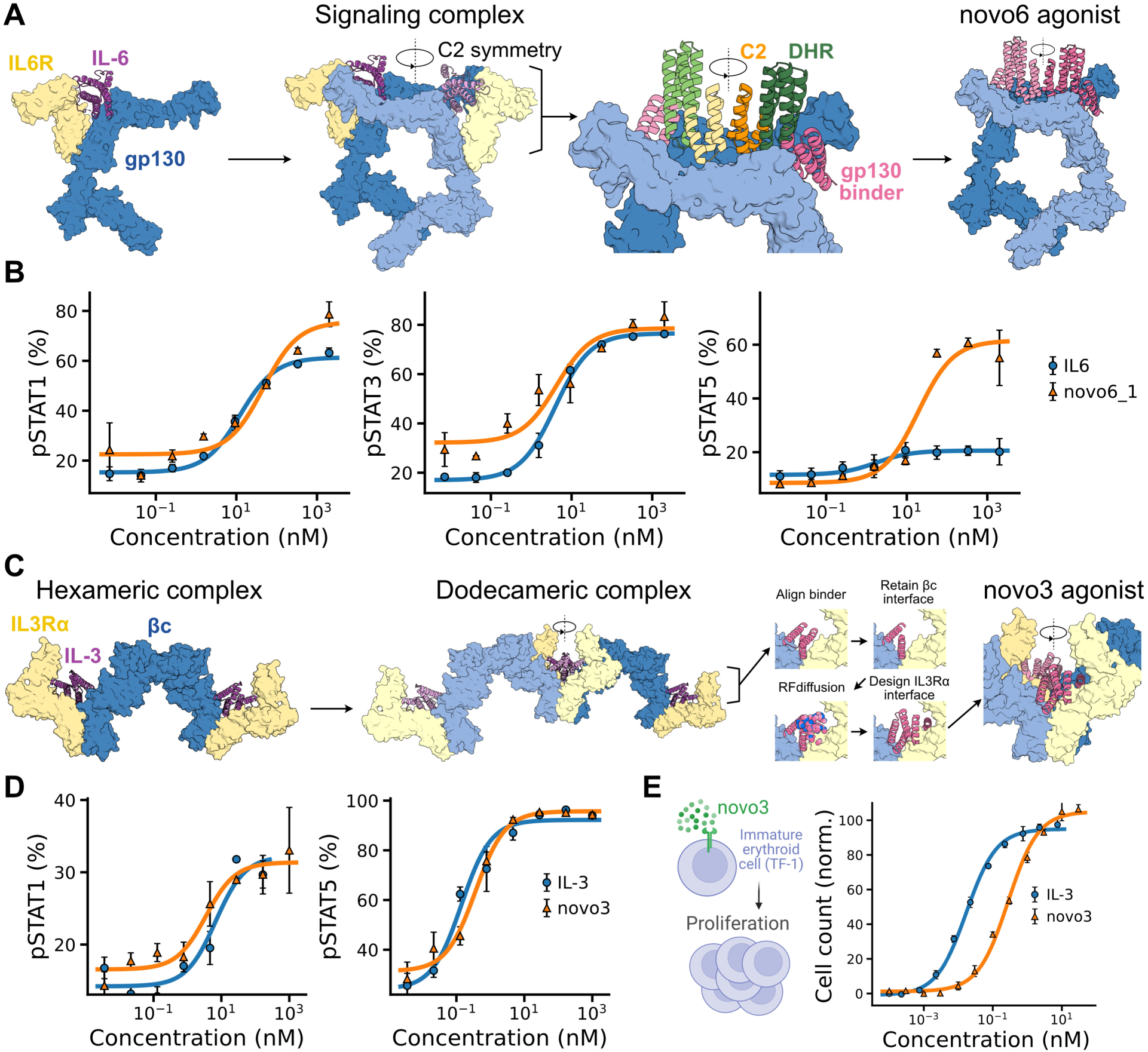
Engineering Minimal and Geometry-Constrained Cytokine Agonists for IL-6 and IL-3. (A) IL-6 signaling complex involving two IL-6 cytokines (pink), two gp130 receptors (blue) with signaling domains, and two IL-6R (yellow) bridging the gap between gp130 chains. Design of symmetric homodimeric cytokines by fusing pre-characterized binders, spacers, and C2s. Minimal IL-6 agonist engaging only gp130 chains. (B) Signaling of IL-6 like *de novo* agonists (novo6_1). (C) IL-3 signaling complex involving two IL-3 cytokines, binding two IL-3Rα receptors and engaging a βc dimer to form hexamers that dimerize and create a dodecameric complex. Design of geometry-constrained agonists starting from a βc binder and creating a custom interface for IL-3Rα using RFdiffusion that satisfies the critical spatial relationships. (D) Signaling activity of a designed *de novo* IL-3-like agonist (novo3). (E) Proliferation of TF-1 cells by the *de novo* designed IL-3-like agonist novo3.

The IL-3 receptor system presented a different challenge, requiring strict preservation of native geometry to support higher-order assembly^48,49^. IL-3 first binds IL-3Rα, which then recruits a homodimer of the β common (βc) subunit to form hexameric and dodecameric signaling complexes through distinct IL-3Rα and βc interfaces (**Figure 5C**)^50,51^. Successful design, therefore, required maintaining the precise spatial relationship between IL-3Rα and βc. To achieve this, we generated a compact *de novo* cytokine that fits within the narrow inter-receptor space. Starting from a pre-designed βc binder, we used RFdiffusion to build a new interface for IL-3Rα while explicitly constraining the design to preserve the native hexameric and dodecameric architecture (**Figures 5C and S9**)^52^. The resulting agonists adopted five-helix bundle topologies; one variant (novo3) mimicked the activity of native IL-3, activating both pSTAT5 and pSTAT1 (**Figure 5D**) and inducing proliferation of IL-3-dependent immature erythroid cells (**Figure 5E**).

Together, these case studies highlight the adaptability of *de novo* cytokine design to multimeric receptor systems. Our framework enables both removal of non-signaling components, as demonstrated with IL-6R, and satisfaction of strict geometric requirements for higher-order complexes, as in IL-3. These strategies expand the design space for cytokine agonists to receptor systems that rely on complex architectures or alternative assembly modes, further generalizing the reach of geometry-based cytokine engineering.

## Discussion

Cytokines are powerful modulators of cellular function, yet their pleiotropy and systemic effects have posed major challenges for therapeutic use, and efforts to engineer biased or selective variants at the protein level remain difficult. Here, we extend the finding that the *geometry* of receptor association, independent of binding affinity, is a powerful axis of control over cytokine signaling, previously engineered only for homodimeric ligands, to the much larger set of ligands inducing association of two distinct receptors, and inducing proximity between receptor pairs not known to be targeted by native ligands. By using *de novo* protein design to precisely position receptor-binding domains in space, we engineered cytokine agonists that tune signaling amplitude, bias pathway activation, and yield novel functions.

Across multiple cytokine families, we find that altering the geometry of receptor engagement can yield biased agonists that selectively modulate functional outputs. In the case of IL-7, a cytokine critical for T-cell homeostasis, we identified designed ligands that decouple and tune pSTAT1 and pSTAT5 signaling, with downstream consequences on gene expression and cell cycle progression. Similarly, for type I interferons, geometric tuning separated the induction of MHC-I and PD-L1, two genes with opposing effects in cancer immunotherapy. The designed novokines maintained strong MHC-I upregulation while reducing PD-L1 overexpression, providing a therapeutic path to enhance tumor antigen presentation without triggering immunosuppressive feedback.

Geometry also provides a route to address long-standing challenges in cytokine therapy. For example, natural IL-10 combines anti-inflammatory effects in monocytes and macrophages with pro-inflammatory IFN-γ induction in CD8+ T cells, complicating its clinical use. We found that dimeric, geometrically constrained IL-10 variants selectively reduced pSTAT1 activation in CD8+ T cells, dampening IFN-γ production while preserving pSTAT3-driven suppression of monocyte activity. These results suggest that IL-10’s opposing activities are not intrinsic properties of the cytokine but emergent features of receptor engagement that can be decoupled through design. Similarly, for type III interferons, whose natural role lies in tissue specificity and reduced systemic toxicity, geometric tuning enabled us to broaden its signaling response.

Finally, we show that geometric design enables the tuning of novel cytokine activities by bringing together receptor pairs not found in nature in a functional signaling orientation. By rigidly scaffolding binders to non-natural receptor combinations, such as IL-2Rβ/IL-10Rβ, yc/IL-10Ra, or IL-21Rα/IFNAR1, we recover signaling that is inaccessible through flexible fusions. These synthetic ligands can activate distinct STAT pathways and reshape T cell phenotypes, potentially eliciting therapeutic effects without the need to engineer cells expressing synthetic receptor pairs. Finally, *de novo* IL-6 and IL-3 designs further show that the same principles can control receptor number and arrangement, a strategy that could extend to other cytokine families to dissect receptor roles and build complexes with tailored composition and function. The ability to rewire signaling in this manner opens new avenues for cytokine-based therapies: rather than engineering native cytokines, one can now design ligands that target specific receptor co-expression profiles and deliver customized intracellular instructions.

Our work positions receptor geometry as an important determinant of cytokine function, on par with receptor identity or binding affinity, and introduces a generalizable platform for creating novel cytokine therapeutics. Beyond immunotherapy, this approach has broad implications for any receptor system governed by proximity-driven activation. As our mechanistic understanding of receptor architecture deepens, and as design tools improve, the ability to precisely sculpt cell signaling is poised to unlock exciting opportunities in programmable immune modulation.

## Acknowledgements

We thank members of the Baker lab for helpful discussions and feedback on the project, L. Goldschmidt and P. Vecchiato for computational infrastructure, and K. VanWormer, H. Nunez-Ortega and R. Ticzon for wet-lab infrastructure. We are grateful to J. Decarreau, S. Cheng, C. Dobbins, and members of the bioassay core for preparing PBMCs and assistance with cell lines, B. Wicky, L. Milles, R. Ragotte, J. Qian, J. Harman and S. Gerben for technical support and protein expression; and D. Lee for RNA-seq guidance. We also thank UW Medicine for access to sequencing instrumentation. Protein visualization was performed using PyMol, and most schematic figures included in this work were created with or modified from BioRender. This work was supported by the Howard Hughes Medical Institute (D.B., J.L.M.), Audacious Project at the Institute for Protein Design (M.E., S.J., D.B.), the Nordstrom Barrier Institute for Protein Design Director’s Fund (M.A., D.B.), the Bill & Melinda Gates Foundation INV-043758 (M.E., W.G., D.B.), and the Microsoft Protein Prediction Research gift. Additional support was provided by the Wu Tsai Protein Innovation Fund (T.S.), and the Damon Runyon Cancer Research Foundation (W.C.). This project was also sponsored by the Fred Hutchinson Cancer Center (DOD Breakthrough W81XWH-20-1-0230), and the National Institutes of Health, including the National Cancer Institute (grants R01CA114536 and R01CA240339). T.S. was supported by the European Molecular Biology Organization (EMBO) via ALTF191-2021. M.H.A. is a Fellow of The Jane Coffin Childs Fund for Medical Research. M.E. acknowledges support from the “La Caixa” Foundation (ID 100010434 under grant no. LCF/BQ/AA19/11720031) and the “Rafael del Pino” Foundation.

## Author Contributions

M.E. and M.H.A. conceived and designed the project, interpreted data, and wrote the manuscript. A.K. and S.J. designed proteins and experiments and analyzed data relevant to Figure 3. Z.C. and W.S.G. conducted experiments for Figure 3. D.M. and S.G.G. performed phenotyping experiments and analyzed the data for Figure 4. W.C. pre-processed RNA-seq data for Figure 1. A.F. and T.S. conceptualized and developed code for the data in Figure 5. T.Y., T.R.H., W.L.K., M.W.P. and A.F.L. conceptualized ideas, designed proteins, performed experiments and interpreted data relevant to Figure 5. D.B. supervised the project. D.B., E.J.W., J.M., and A.F.L. contributed to data review and interpretation. All authors participated in editing the manuscript.

## Data and Code Availability

All scripts involved in the design of rigid novokines will be made available upon final publication. All raw data and analysis associated with this study not currently present within the manuscript will be provided within supplementary information files.

## Declaration of Interests

D.B., M.E., M.H.A., and A.K. are inventors of a provisional patent application submitted by the University of Washington for the designed novokines. E.J.W. is a member of the Parker Institute for Cancer Immunotherapy which supported this study. E.J.W. is an advisor for Arpelos Biosciences, Arsenal Biosciences, Coherus, Danger Bio, IpiNovyx, New Limit, Marengo, Pluto Immunotherapeutics, Related Sciences, Santa Ana Bio, and Synthekine. E.J.W. is a founder of Arpelos Biosciences, Danger Bio, and Arsenal Biosciences. E.J.W. holds stock in Coherus. All other authors declare no competing interests.

## Supplementary Figures

**Figure S1.**
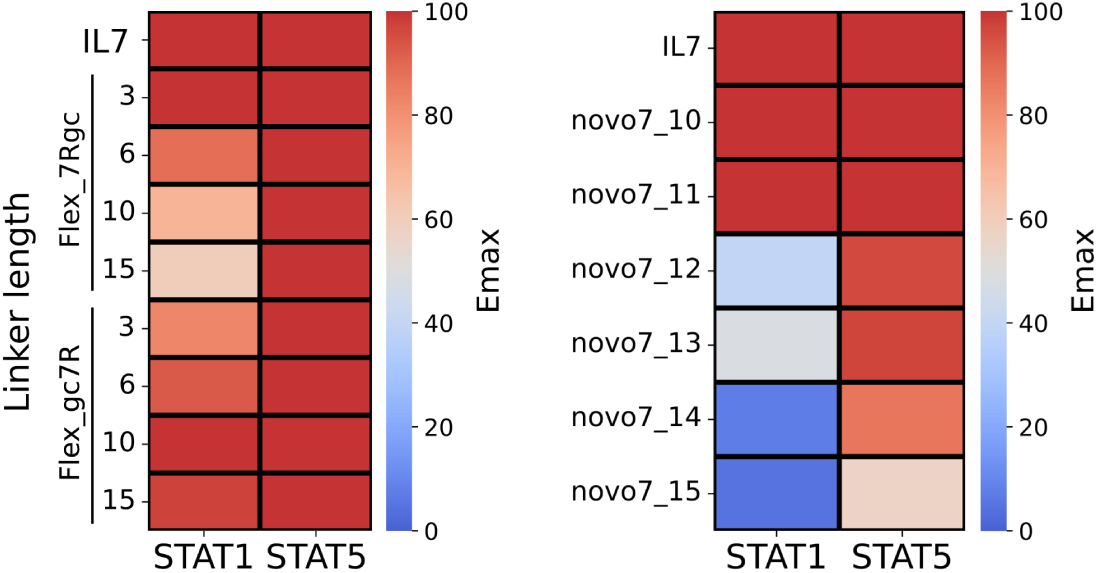
Reconstructing IL-7 signaling by flexibly fusing *de novo* binders. (left), Signaling strength (Emax) for STAT1 and STAT5 for designed IL-7Rα and γc binder flexible fusions. The orientation of the fusion (IL-7Rα-γc or γc-IL-7Rα) and length in amino acids of the flexible (GGS) linker is specified for each agonist. (right), Signaling strength for designed rigid novokines.

**Figure S2.**
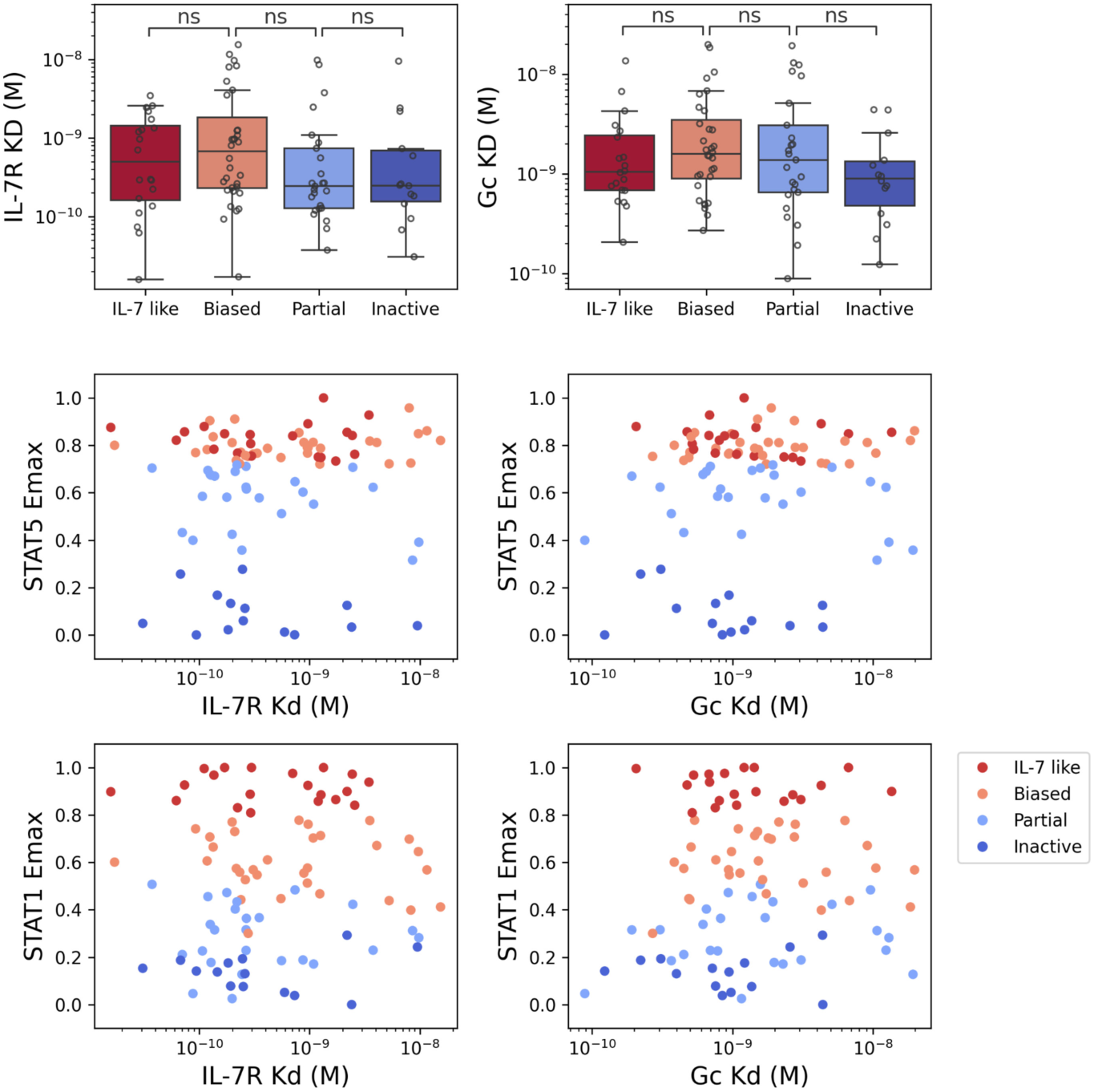
Relation between affinity (K_D_) and signaling strength (E_max_) for designed IL-7 novokines. (top), Affinity of novo7 novokines grouped by their signaling pattern. n.s.= Not significant (p-value>0.05). (middle), scatterplot of affinities for each receptor and relation to pSTAT5 Emax. (bottom), scatterplot of affinities for each receptor and relation to pSTAT1 Emax.

**Figure S3.**
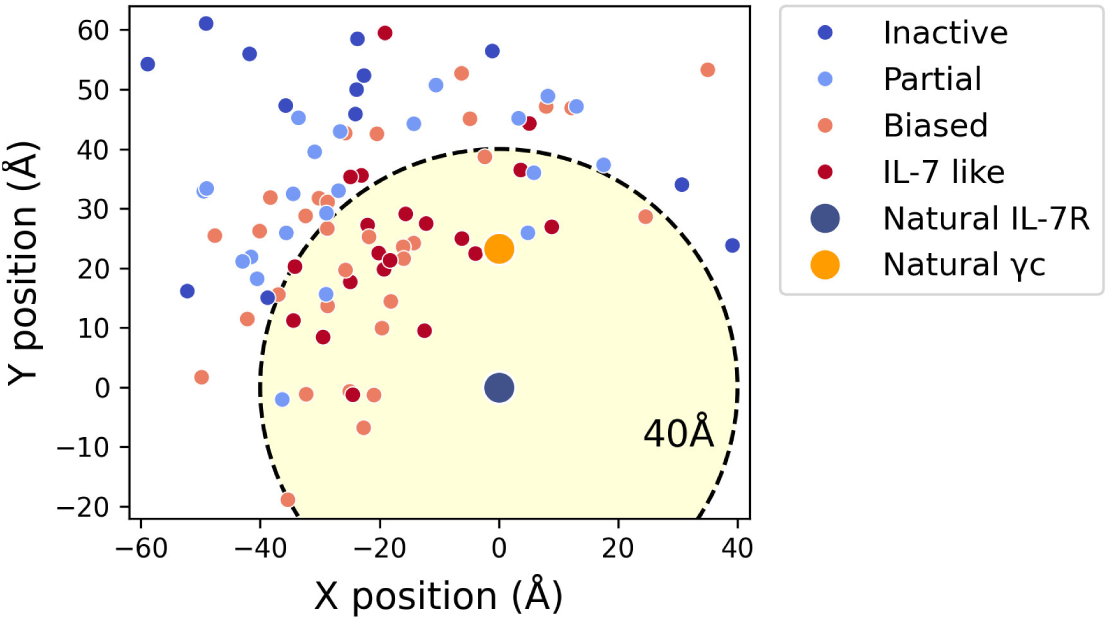
X-Y positions of receptor termini for novo7 novokines. Membrane-proximal receptor termini positions in a X-Y plane for designed agonists, colored by signaling pattern, all aligned using the IL-7Rα receptor as reference, and including the position of the γc in the natural IL-7 complex.

**Figure S4.**
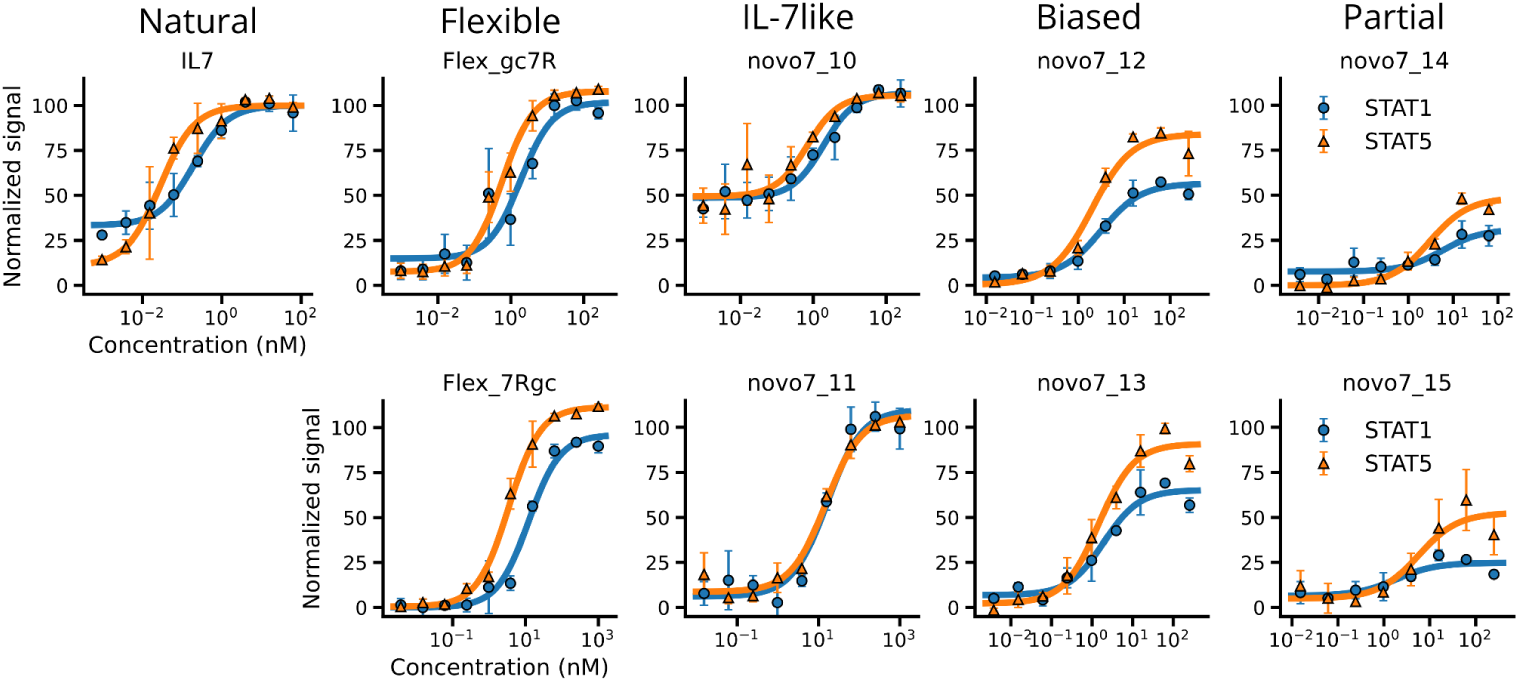
Dose curve for novo7 novokines analyzed by RNA-seq. pSTAT1 (blue) and pSTAT5 (orange) signaling for novo7 agonists analyzed in RNA-seq, organized by signaling pattern, including the natural IL-7 cytokine and flexible fusions between IL-7Rα and γc binders.

**Figure S5.**
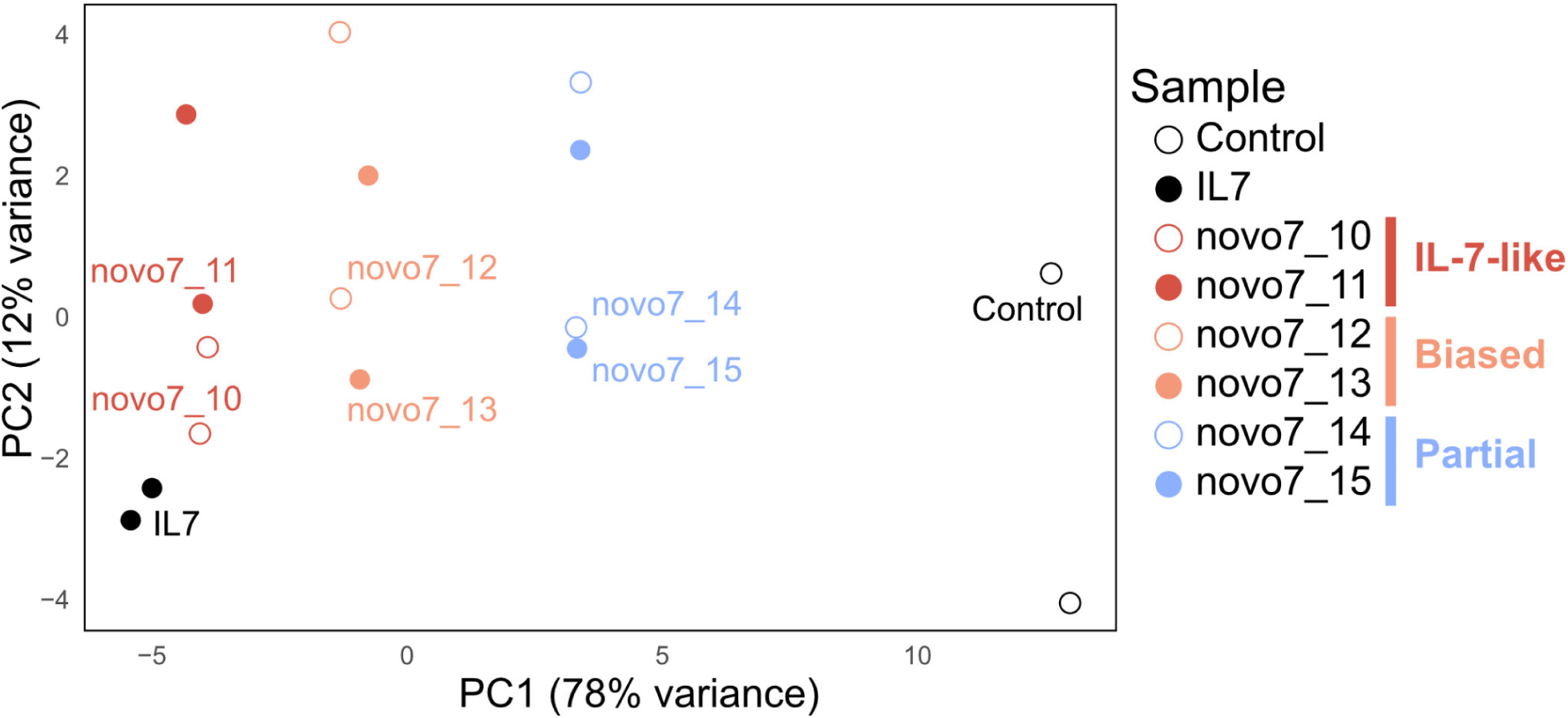
Principal component analysis (PCA) of RNA-seq samples colored and shaped by sample identity. Each point represents an individual RNA-seq replicate, with colors indicating functional classifications (e.g., IL-7-like, biased, partial), and point shapes distinguishing the two samples within each group. Each sample has two biological replicates. Percent variance explained by PC1 and PC2 is indicated on the axes. PC1 separates samples based on functional phenotype, while PC2, accounting for a smaller proportion of variance, captures intra-group replicate variability.

**Figure S6.**
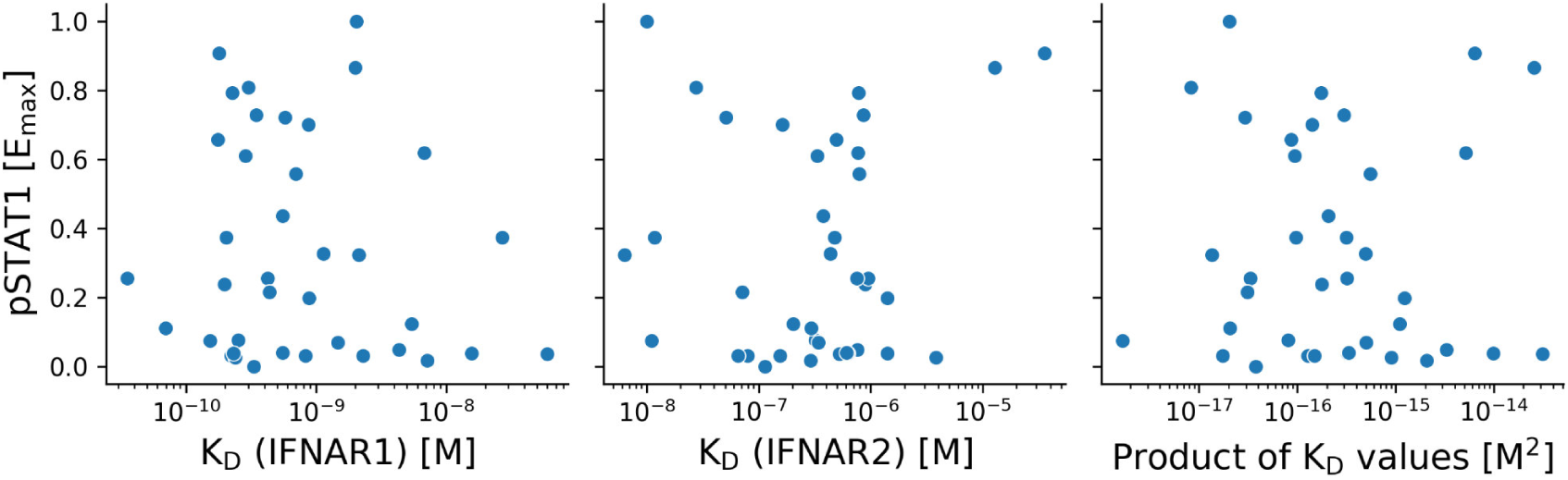
Relation between affinity (K_D_) and signaling strength (E_max_) for designed type I IFN novokines. Scatterplots between pSTAT1 Emax normalized with respect to natural IFNa1, and relation to affinity measured by SPR for IFNAR1 (left), IFNAR2 (center) or the product between both (right).

**Figure S7.**
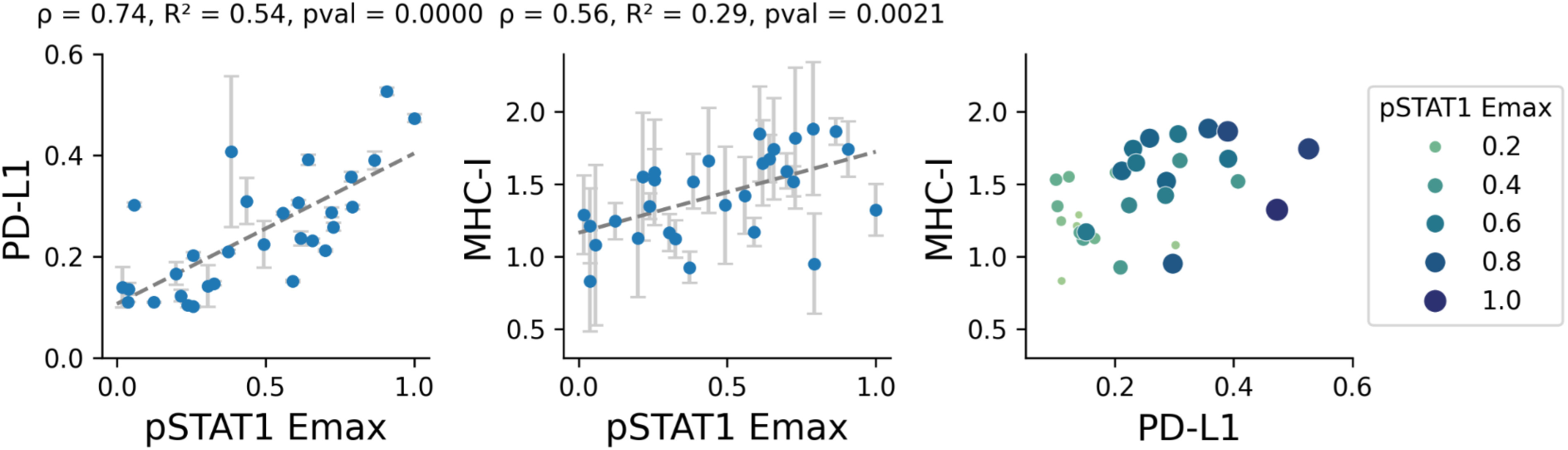
Scatterplots of PD-L1 and MHC-I expression in relation to pSTAT1 signaling strength. Surface expression levels of PD-L1 (left) and MHC-I (center), normalized to natural IFNα1, are plotted against the maximal pSTAT1 signaling response (Emax) elicited by each designed novoIFNα ligand. Dotted lines indicate linear regression fits, with Spearman’s correlation coefficient (ρ), R², and permutation-derived p-values shown above each plot. The right panel displays a scatterplot of MHC-I versus PD-L1 expression, with point size reflecting the pSTAT1 Emax for each novoIFNα ligand.

**Figure S8.**
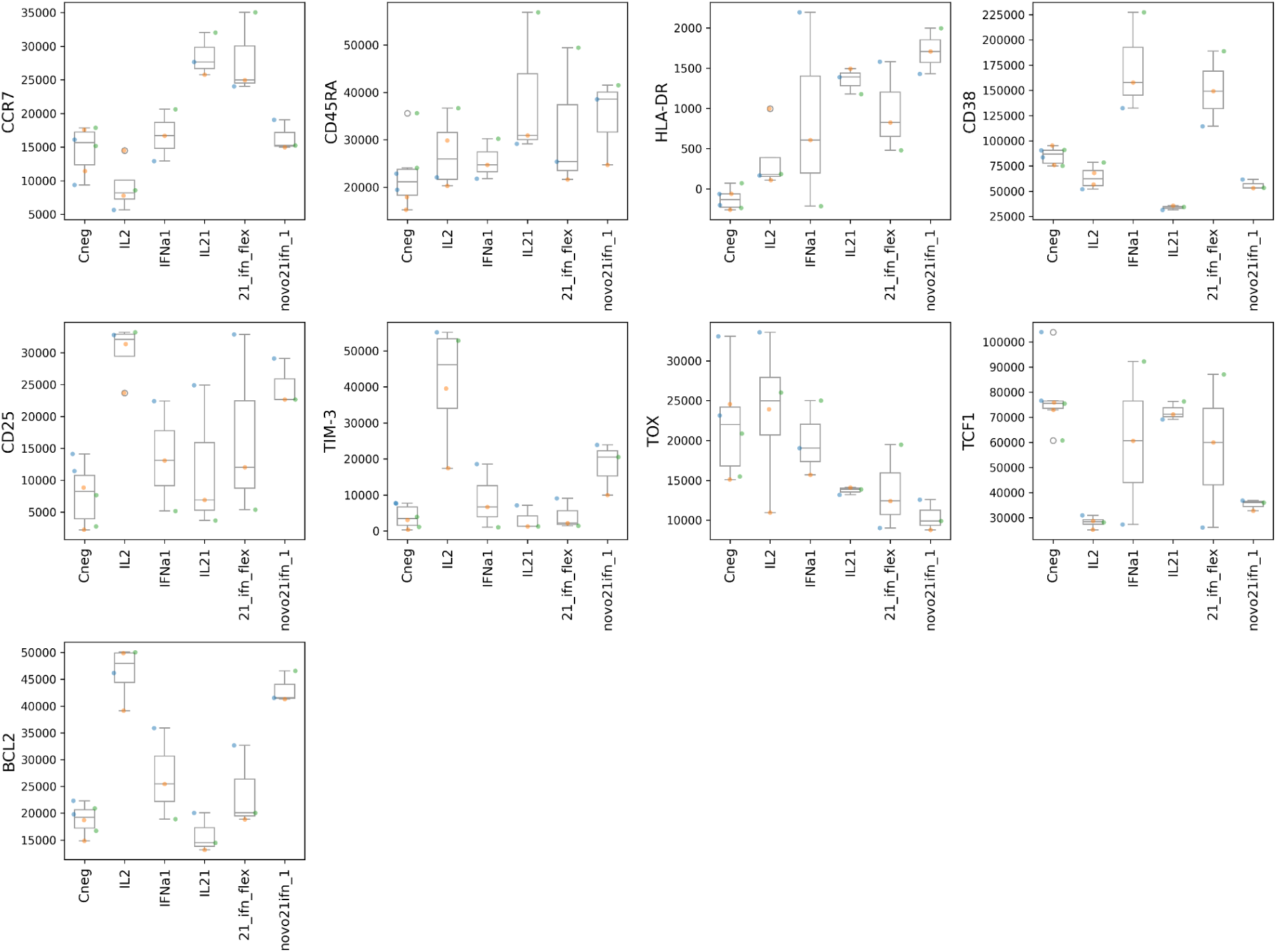
Box plots of selected markers in the naive CD8^+^ T cell differentiation assay. Each color indicates a different donor. Units are directly the values obtained from flow cytometry.

**Figure S9.**
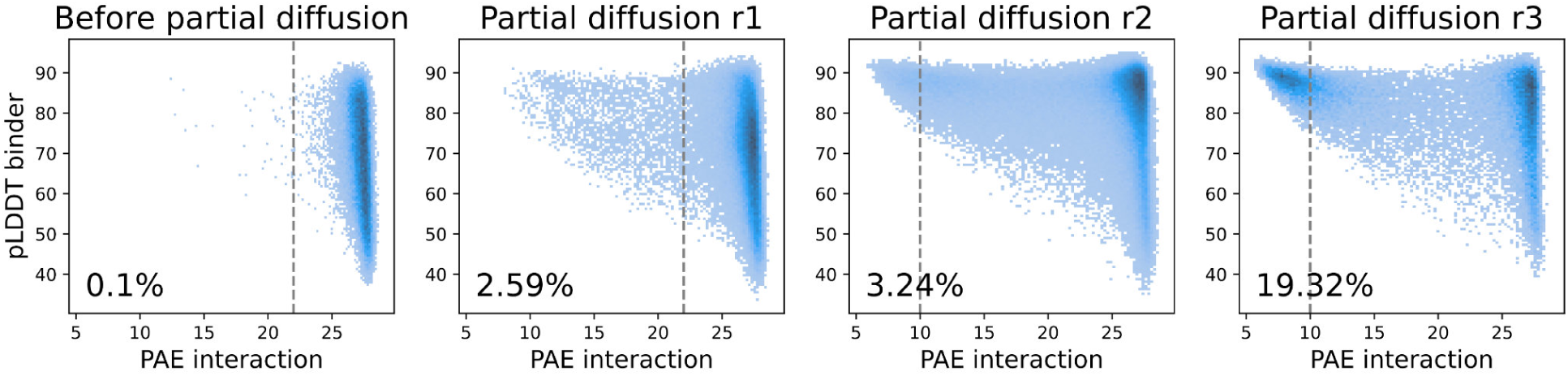
Improvement of binding affinity for IL-3Rα through multiple rounds of partial diffusion. Scatterplots between AlphaFold2’s accuracy of prediction (pLDDT) and strength of interaction (PAE) for designed novo3 novokines across rounds of partial diffusions. The percentage of designs below a PAE threshold is shown for comparison. Final designs were below PAE<10 after the 3rd round of diffusion.

## Methods

### Design of *de novo* binders

We designed binders against human receptors using reference structures including IL-10Rβ (PDB ID: 6X93) and IFN-λ (PDB ID: 5T5W)^33,53^. The design process is similar to the one described previously by Watson et al.^54^. Briefly, RFdiffusion was used to generate protein backbones through iterative denoising, conditioned to target specific hot spot residues within the receptor binding interface that would antagonize the natural cytokine. ProteinMPNN was then used to design sequences for the generated backbones and a modified AlphaFold2 (AF2) used to filter for high pLDDT (confidence in predicted structure) and low PAE for binding likelihood^55,56^. Designs were screened for binding to their target using Surface Plasma Resonance (SPR, described below) and hits were further optimized through rounds of partial diffusion where the interface is allowed to perturb slightly and the sequence is redesigned with ProteinMPNN and filtered with AF2^57,58^.

### Initial structures and previously described binders

For each natural receptor pair, we used either available complex structures or homology models based on structurally related cytokines as the initial reference for geometric sampling. In the case of IL-7, no structure of the complex with the common gamma chain (γc) receptor is available. Therefore, we used the structure of a previously published IL-7R binder (PDB ID: 7OPB) and overlaid it with structurally homologous cytokines, including TSLP (PDB ID: 4NN5), IL-2 (2ERJ), and IL-4 (3BPL)^17^. To engage γc, we incorporated the γc binder described in the companion paper by Abedi et al.^18^.

For type I interferons, we started from the IFNAR1-IFNAR2-IFNα2 complex structure (PDB ID: 3SE3)^26^. To model the complete extracellular geometry, a full-length IFNAR1 structure predicted with AF2 that includes the membrane-proximal domain was overlaid^59^. IFNAR1 and IFNAR2 binders described in the companion paper by Abedi et al. were used to design the novokine^18^. For IL-10, we used the structure of the IL-10 signaling complex (PDB ID: 6X93) in combination with a previously described IL-10Rα binder and a newly developed IL-10Rβ binder^33,55^. For IFN-λ, we started with a high-affinity IFN-λ variant bound to its receptors IFN-λR1 and IL-10Rβ (PDB ID: 5T5W), incorporating the newly designed binders for both IFN-λR1 and IL-10Rβ in the modeling process^53^.

For non-natural cytokine receptor pairs (IFNAR1-IL-2Rβ, IL-10Rβ-IL-2Rβ, IL-10Rα-γc, and IL-21Rα-IFNAR1), we estimated an initial geometry by positioning the membrane-proximal termini of the receptors in close proximity while avoiding steric clashes between any part of the receptors or binders. For these cases, in addition to the previously mentioned binders, we used the IL2Rβ binder described in the companion paper by Abedi et al. and a previously designed IL-21R binder^18,60^.

### Design of rigid agonists

The reference complex structure for each cytokine-receptor system was used to guide geometric sampling. Receptor-binder complexes were centered in a standardized coordinate space, with the origin placed at the midpoint between the two receptor centers. The Z-axis was defined as perpendicular to the plane of the cell membrane; the X-axis aligned with the vector connecting the membrane-proximal domains of the two receptors; and the Y-axis was defined as orthogonal to both X and Z. This alignment enabled systematic parameterization of sampling and rotation in biologically meaningful directions. A custom PyMOL script was used to sample thousands of geometries by fixing one receptor and systematically translating or rotating the other along specific axes. Rotations along the Z-axis were extensively sampled, as well as translations to increase inter-receptor distance or reorient the complex to position intracellular domains in proximity but at variable angles. The sampling followed a Gaussian distribution centered on the natural geometry, enabling focused yet broad spatial coverage. The script also included clash-filtering steps to eliminate binder-receptor conformations with significant steric clashes, ensuring only physically plausible configurations were passed forward for design.

Next, the two binders were fused into a rigid, single-chain protein using RFdiffusion, treating each binder as a contig. Variable-length scaffolds were generated based on the distance between the N- and C-termini of the binders. Both fusion orientations were attempted, i.e. connecting binder A’s C-terminus to binder B’s N-terminus and vice versa. For the IFN-λ system, guiding helices were incorporated to promote compact, rigid fusions. After diffusion, the receptors were aligned back to the outputs to filter out any scaffolds that clash with them. The original binder sequences were then threaded onto the designs to preserve as much of the original sequence as possible. Residues in or contacting the new scaffold (within 5 Å) were redesigned using ProteinMPNN, excluding residues directly involved in receptor interfaces. Designs were filtered using AF2 monomer predictions to assess model fidelity. Designs with low RMSD to the input model, high pLDDT scores (AF2 confidence), and sampling a diverse range of receptor geometries were prioritized for experimental testing.

### Design of a minimal gp130 signaling complex

The crystal structure of the IL-6 hexameric signaling complex (PDB ID: 1P9M), comprising two IL-6 agonists, two gp130 receptors, and two IL-6R, was used as a reference to overlay the full ectodomain of gp130 (PDB ID: 3L5H) so the model includes the membrane-proximal domains^45,61^. Using Pymol, the complex was centered and aligned such that the Z-axis of the coordinate system was aligned with the C2 symmetry axis of the hexameric complex, which is oriented perpendicular to the membrane. A previously described gp130 binder was aligned to both gp130 chains^62^, and multiple geometries were generated by manually sampling binder positions to explore different orientations and distances of the membrane-proximal domains without receptor clashes.

The gp130 agonists were assembled using the WORMS software, which facilitates helical fusion by connecting protein segments through similar helices^47^. First, the C-terminus of the binder was extended by 10 residues using Rosetta’s Blueprint Builder^63^, providing an anchor for helix fusion. A database was generated containing binder geometries, designed helical repeats (DHRs), and *de novo* C2 homodimers^64^. The WORMS protocol connected the binder to a DHR spacer, which was then fused to the C2 scaffold centered at the Z-axis to form a homodimeric agonist, ensuring no clashes with the receptors or other components. Geometries were filtered to maintain the correct binder position. Subsequently, junctions between components were redesigned using Rosetta FastDesign, ensuring that no residues of the binder or C2 interface were modified. The final designs were predicted using AF2 for both monomers and homodimers. Structures with high pLDDT scores and low RMSD to the design models, representing diverse geometries, were selected for further analysis, yielding a subset of 48 for ordering.

### Design of an IL-3-like agonist

The structure of β common (βc) in complex with IL-3 and IL-3Rα (PDB ID: 6NMY) was used as a reference for designing the novo3 agonist^52^. The βc binder, described in Abedi et al.^18^, was aligned with the βc receptor, retaining only the two helices that interact with the receptor. The IL-3Rα interface design followed the protocol from Watson et al.^54^, with modifications to incorporate the pre-existing helices. Specifically, the helices were integrated into two separate contigs, placed at either the N-terminus, C-terminus, or mid-sequence, with all possible permutations tested between the generated structure and each helix. To speed up RFdiffusion runs, the IL-3Rα was cropped to include only residues in proximity to the relevant binding domain, and the βc chain was excluded from the RFdiffusion input. Additionally, all residues of the βc interface within the two retained helices were mutated to glutamic acid, preventing the generated protein from interacting there and ensuring that the Bc interface remained unblocked. Hotspot residues within IL-3Rα (A72, V201, and Y279) were selected for targeted interaction.

Initial filtering of RFdiffusion output removed designs with the lowest number of residues in contact (<5 Å) with IL-3Rα, as these were unlikely to form a stable interface. Promising designs were selected for sequence design. The original sequence was reintroduced, reverting the glutamic acid mutations, and ProteinMPNN was employed to design the generated regions and any residues that interacted with it, but excluding any on the pre-existing βc interface. The full IL-3Rα receptor was reincorporated into the design, and binding predictions were made using a modified AF2 protocol for binder prediction as previously described^55^. Designs that exhibited high pLDDT, low PAE, and low RMSD were prioritized. Three rounds of partial diffusion of the IL-3Rα interface were performed until enough designs with PAE<10 were identified (Figure S9). In the final round, designs were validated in the context of both βc and IL-3Rα structures to ensure compatibility and minimize clashes within the complex.

### Protein expression of agonists

For protein expression, we followed a previously reported protocol with some modifications^65,66^. Golden Gate subcloning reactions (1 µL) were transformed into BL21(DE3) cells, which were outgrown for 1 hour and split into four 96-deep well plates containing ∼1 mL auto-induction media per well. After 20–24 hours, cells were harvested and lysed. The insoluble fractions were resuspended in 6M guanidine hydrochloride and applied to 50uL Ni-NTA resin in 96-well fritted plates. Proteins were gradually refolded through four washes with 2-fold decreasing guanidine concentrations, followed by addition of soluble fractions. Resins were washed with a CHAPS-containing buffer to remove endotoxin, with standard Tris buffer to remove CHAPS excess, and proteins were eluted in 200 µL Tris buffer with 300 mM imidazole. Eluates were sterile-filtered (0.22 µm) prior to size exclusion chromatography (SEC).

Proteins were analyzed on an AKTA FPLC with an autosampler and either a Superdex 75 Increase 5/150 GL column (Cytiva Cat# 29148722) or a Superdex 200 Increase 5/150 GL column (Cytiva Cat# 28990945). The SEC column was connected directly from the autosampler to the UV detector for improved resolution. Fractions of 0.25 mL were collected, and selected fractions were pooled using an OT-2 robot (Opentrons Cat# 999-00111) for further analysis.

### Experimental Screening of Binders

Binding affinity to target receptors was measured using a Biacore 8K with a Protein A chip (Cytiva #29127556) at a target concentration of 200ng/mL. Binding traces were collected by running single-cycle kinetics at increased binder concentrations (four series with 8-fold titration starting at 1000 nM or 1000nM), with binders and targets diluted in HEPES buffer (Cytiva #BR100669).

### Cell Lines and PBMCs

For this study, the following cell lines were used: wild-type NALM-6, THP-1, TF-1, A379, and A549 (ATCC). All suspension cell lines (NALM-6, THP-1, TF-1) were cultured in RPMI-1640 Medium with 1% GlutaMAX (Gibco Cat# 61870036), supplemented with 10% fetal bovine serum (Life Technologies Cat# A4766801) and 1% penicillin-streptomycin (Gibco Cat# 15140122). TF-1 cells were additionally supplemented with 2 ng/mL of recombinant human GM-CSF (Biotechne Cat# 7954-Gm) to support their growth. Adherent cell lines (A379, A549) were cultured in DMEM Medium with 1% GlutaMAX (Gibco Cat# 10566016), supplemented with 10% fetal bovine serum (Life Technologies Cat# A4766801) and 1% penicillin-streptomycin (Gibco Cat# 15140122). The cells were passed every 3 days to avoid over-confluency and maintained at 37°C in a humidified incubator with 5% CO2. Peripheral Blood Mononuclear Cells (PBMCs) were obtained from Bloodworks Seattle, thawed, washed with CTL Anti-Aggregate Wash (ImmunoSpot Cat# CTL-AA-010), and either rested for one hour before experimentation or subjected to a 3-hour starvation period prior to assays if required.

### Intracellular Signaling and Surface Expression Assays

Intracellular phospho-staining assays were performed using a unified protocol. Cells were stimulated with the relevant ligand for 15 minutes, then fixed with BD Cytofix Fixation Buffer (BD #554655) for 10 minutes and permeabilized with BD Phosflow Perm Buffer III (BD #558050). Cells were stained with the following phospho-specific antibodies: Alexa Fluor 647 Mouse Anti-STAT1 (pY701) (BD #562070), PE Mouse Anti-STAT5 (pY694) (BD #562077), and Alexa Fluor 488 Mouse Anti-STAT3 (pY705) (BD #557814). For IL-10 experiments, STAT1 and STAT3 staining was performed together with surface markers for CD8^+^ T cells (PE anti-human CD8, Miltenyi Biotec #130-113-158) and monocytes (Pacific Blue anti-human CD14, BioLegend #367122). For IFN-λ and novoIFNλ stimulation, the epithelial cell line A549 was used since the IFN-λR1 is not present in PBMCs. For experiments, adherent A549 cells were gently detached using enzyme-free Cell Dissociation Buffer (Gibco #13151014) for 30 minutes prior to stimulation. Phospho-signal quantification was performed on an Attune NxT Flow Cytometer. Data analysis and dose-response curve fitting were conducted using custom Python notebooks based on FlowKit^67^.

To study PD-L1 and MHC-I expression in response to type I IFNs or designed novoIFNα, A379 cells were incubated overnight with 100nM of IFNa1 or a designed novoIFNα agonist, followed by surface staining with PD-L1 and MHC-I antibodies and analyzed on flow cytometry. To assess the significance of the observed correlations, we performed a permutation test (10,000 permutations) by randomly shuffling the response variable (PD-L1 or MHC-I) to generate a null distribution of R² values and computed a p-value as the fraction of null R² ≥ observed R².

### Cell proliferation and Differentiation Assays

For immature erythroid cell proliferation assay, erythroleukemic TF-1 cells were starved overnight in RPMI medium + 10% FBS without GM-CSF. The following day, 5,000 cells/well were seeded in 96-well plates and stimulated with the indicated concentrations of IL-3 or novo3. After 72h, cell proliferation was measured using CellTiter 96® Aqueous 703 (Promega Cat # G3581) following the manufacturers protocol.

For the naive CD8^+^ T-cell differentiation assay, naive CD8^+^ T cells were isolated from PBMCs by negative magnetic bead enrichment using EasySep™ Human Naïve CD8+ T Cell Isolation Kit following the manufacturers protocol (StemCell Technologies Cat# 19258). Purified naive CD8^+^ T cells were then treated with cell trace violet, and incubated for 4 days with CD3/CD28 beads (Gibco Cat# 111.31D) and supplemented with a natural cytokine (IL-2, IL-21, or IFNa1) or a synthetic novokine (novo21ifn_1, or 21_ifn_flex). Experiments were performed in triplicate using 3 different PBMC donors. After 4 days, cells were harvested, stained with a panel of surface antibodies for selected markers, and ran on flow cytometry.

For T cell proliferation assays, CD4^+^ and CD8^+^ T cells were isolated from PBMCs following the manufacturers protocol (StemCell Technologies Cat# 17952 and Cat# 100-0710). Purified T cells were then incubated with CD3/CD28 beads and IL-2 for 2 days. On the third day, the beads were removed, and the cells were distributed into a 384-well plate. Designed ligands were added to each well in triplicate, with ligands replenished every 2-4 days. Cells were passed based on confluency, with the test ligands refreshed accordingly. At passing steps after 10, 14 and 18 days, leftover cells from passing were counted using flow cytometry.

### Cytokine Secretion Assays

For TNFα measurement in the context of bulk PBMCs, PBMCs were thawed as described above and plated in a 96 well U-bottom plate at 100,000 cells/well, followed by bacterial LPS stimulation at 1ng/ml in the absence or presence of 10nM IL-10 or novo10 designs for 24 hours. The following day TNFα secretion was measured in the cell supernatants using a commercially available ELISA kit (R&D Human TNFα ELISA Quantikine kit, Biotechne Cat# DTA00D).

For IFNy measurements in the context of CD8^+^ T cells, CD8^+^ T cells were isolated from bulk PBMCs as described above. Cells were then activated and expanded for 3 days using CD3/CD28 beads in RPMI media supplemented with IL-2. On day 3, beads were removed and cells were resuspended in fresh media without IL-2 for 24 hours. Cells were then seeded in a 96 well U-bottom plate at 100,000 cells/well and stimulated with 10nM IL-10 or novo10 designs for 72 hours. Following this, cells were restimulated with anti-CD3 OKT3 antibody (ThermoFischer Cat# 14-0037-82) for 4 hours. Supernatants were collected and IFNy levels were measured using a commercially available ELISA kit (R&D Quantikine Human IFNy kit, Biotechne Cat# DIF50C)^33^.

### RNA-seq and GO term annotation

CD4+ T cells were isolated and stimulated overnight with IL-7 or designed novo7 agonists. Total RNA was extracted using standard methods and used to prepare sequencing libraries, which were sequenced on an Illumina platform. Raw reads were quality-checked, trimmed for adapters and low-quality bases, and aligned to the reference genome using standard pipelines. Gene-level counts were quantified, and differential expression analysis was performed with DESeq2, averaging over two biological replicates per condition and comparing each ligand to an unstimulated control. The top 25 most significantly upregulated or downregulated genes for each ligand were visualized in heatmaps. Functional enrichment analysis was performed using Gene Ontology (GO) terms, focusing on biological processes relevant to T-cell signaling and proliferation. To assess global transcriptional differences, principal component analysis (PCA) was conducted on variance-stabilized expression values obtained in DESeq2.

